# Network Modeling Predicts Personalized Gene Expression and Drug Responses in Valve Myofibroblasts Cultured with Patient Sera

**DOI:** 10.1101/2021.09.04.458984

**Authors:** Jesse D. Rogers, Brian A. Aguado, Kelsey M. Watts, Kristi S. Anseth, William J. Richardson

**Affiliations:** Bioengineering Department, Clemson University, Clemson, SC; Center for Computational Toxicology and Exposure, U.S. Environmental Protection Agency, Durham, NC; Chemical and Biological Engineering Department, BioFrontiers Institute, University of Colorado, Boulder, CO; Bioengineering Department, University of California San Diego, La Jolla, CA; Sanford Consortium for Regenerative Medicine, La Jolla, CA

**Keywords:** Personalized medicine, signaling network, heart valve, myofibroblast, computational model

## Abstract

Aortic valve stenosis (AVS) patients experience pathogenic valve leaflet stiffening due to excessive extracellular matrix (ECM) remodeling. Numerous microenvironmental cues influence pathogenic expression of ECM remodeling genes in tissue-resident valvular myofibroblasts, and the regulation of complex myofibroblast signaling networks depends on patient-specific extracellular factors. Here, we combined a manually curated myofibroblast signaling network with a data-driven transcription factor network to predict patient-specific myofibroblast gene expression signatures and drug responses. Using transcriptomic data from myofibroblasts cultured with AVS patient sera, we produced a large-scale, logic-gated differential equation model in which 11 biochemical and biomechanical signals are transduced via a network of 334 signaling and transcription reactions to accurately predict the expression of 27 fibrosis-related genes. Correlations were found between personalized model-predicted gene expression and AVS patient echocardiography data, suggesting links between fibrosis-related signaling and patient-specific AVS severity. Further, global network perturbation analyses revealed signaling molecules with the most influence over network-wide activity including endothelin 1 (ET1), interleukin 6 (IL6), and transforming growth factor β (TGFβ) along with downstream mediators c-Jun N-terminal kinase (JNK), signal transducer and activator of transcription (STAT), and reactive oxygen species (ROS). Lastly, we performed virtual drug screening to identify patient-specific drug responses, which were experimentally validated via fibrotic gene expression measurements in VICs cultured with AVS patient sera and treated with or without bosentan - a clinically approved ET1 receptor inhibitor. In sum, our work advances the ability of computational approaches to provide a mechanistic basis for clinical decisions including patient stratification and personalized drug screening.

## Introduction

Valvular and myocardial fibrosis progression is a major obstacle for the treatment of patients suffering from aortic valve stenosis (AVS). AVS is characterized by thickening of aortic valve leaflets via progressive extracellular matrix (ECM) accumulation and calcification, reducing blood flow through the valve and leading to compensatory cardiac hypertrophy, myocardial fibrosis, and eventually heart failure^1^. Minimally invasive transcatheter aortic valve replacements (TAVR) serve as the current gold standard for treating severe AVS patients with high surgical risk^2^. Unfortunately, interstitial fibrosis prior to surgery is not necessarily reversed after valve replacement, and high degrees of fibrosis are associated with reductions in long-term survival and cardiac function^3–5^. Therefore, preventing fibrosis during initial AVS progression remains crucial for preserving heart function and preventing the late onset of heart failure. Identifying appropriate therapeutic compounds for this task remains a significant clinical need^6^.

As mediators of ECM remodeling, valvular myofibroblasts play key roles during AVS progression, synthesizing matrix proteins, proteases such as matrix metalloproteinases (MMPs) and other regulatory matricellular proteins that orchestrate scar formation and increase tissue stiffness^7^. Valvular interstitial cells (VICs) undergo activation to a myofibroblast phenotype in response to an influx of inflammatory cytokines, growth factors, and hormonal peptides, as well as changes in local biomechanical cues including pathological stiffness and tensile loading^8–10^. These extracellular cues are transduced through a complex network of regulation in which multiple intracellular signaling pathways interact via crosstalk and feedback mechanisms to dictate overall cell responses^11^, thus further complicating efforts to understand how myofibroblasts respond within the valve microenvironment. While previous studies have shown that myofibroblasts can undergo deactivation in response to serum biomarkers after TAVR, macrophage-secreted inflammatory cues, and local mechanical stiffness^12–16^, the complexities of myofibroblast signal transduction among a diverse microenvironment complicates efforts to control cell deactivation.

To account for the complex system of myofibroblast intracellular signaling, we employed a computational model in which environmental cues are transduced via a network of intracellular signaling and transcriptional interactions to predict changes in protein expression. Similar approaches have been used to model various cardiovascular and systemic diseases and successfully generate insight into influential mechanisms within this broad biological scope^17–21^. In our current work, we adapted these approaches to model myofibroblast responses to patient-specific serum biomarker levels before and after TAVR by combining a published network of fibroblast signaling with a new, data-driven network of myofibroblast transcriptional regulation. We developed network reaction topology and fit reaction parameters using VIC RNA-sequencing data under various patient-matched serum conditions, then tested the correlations of personalized model predictions to clinically measured, patient-specific AVS severity. In addition, we conducted network perturbation analyses and patient-specific drug target screens, followed by in vitro validation experiments in order to test the model’s ability to predict variable drug responses across diverse patient sera contexts.

## Results

### Development of Myofibroblast Transcriptional Network

Using a published dataset of valvular myofibroblast gene expression in response to pre- and post-TAVR patient sera^12^, we developed a new model of myofibroblast transcriptional regulation using established gene regulatory network inference algorithms. Upon initial topology inference and implementing several filtering steps to ensure sufficient experimental evidence and relation to fibrosis-related gene expression (Figure 1A), our transcriptional network represents 10 transcription factors (TFs) activated by intracellular signaling, 27 intermediate TFs, and 18 target genes related to ECM turnover or autocrine feedback (Figure S1). This network inference strategy identified mechanisms mediating the expression of structural matrix proteins, matricellular proteins, and remodeling proteins related to collagen processing, crosslinking, and turnover. Additionally, this data-driven approach identified several intermediate TFs known to regulate cardiac myofibroblast behavior during fibrosis, including SOX9^22,23^, CCND1^24^, ATF3^25,26^, and NOTCH1^27^, suggesting agreement between inferred topology and previous experimental findings.

**Figure 1.**
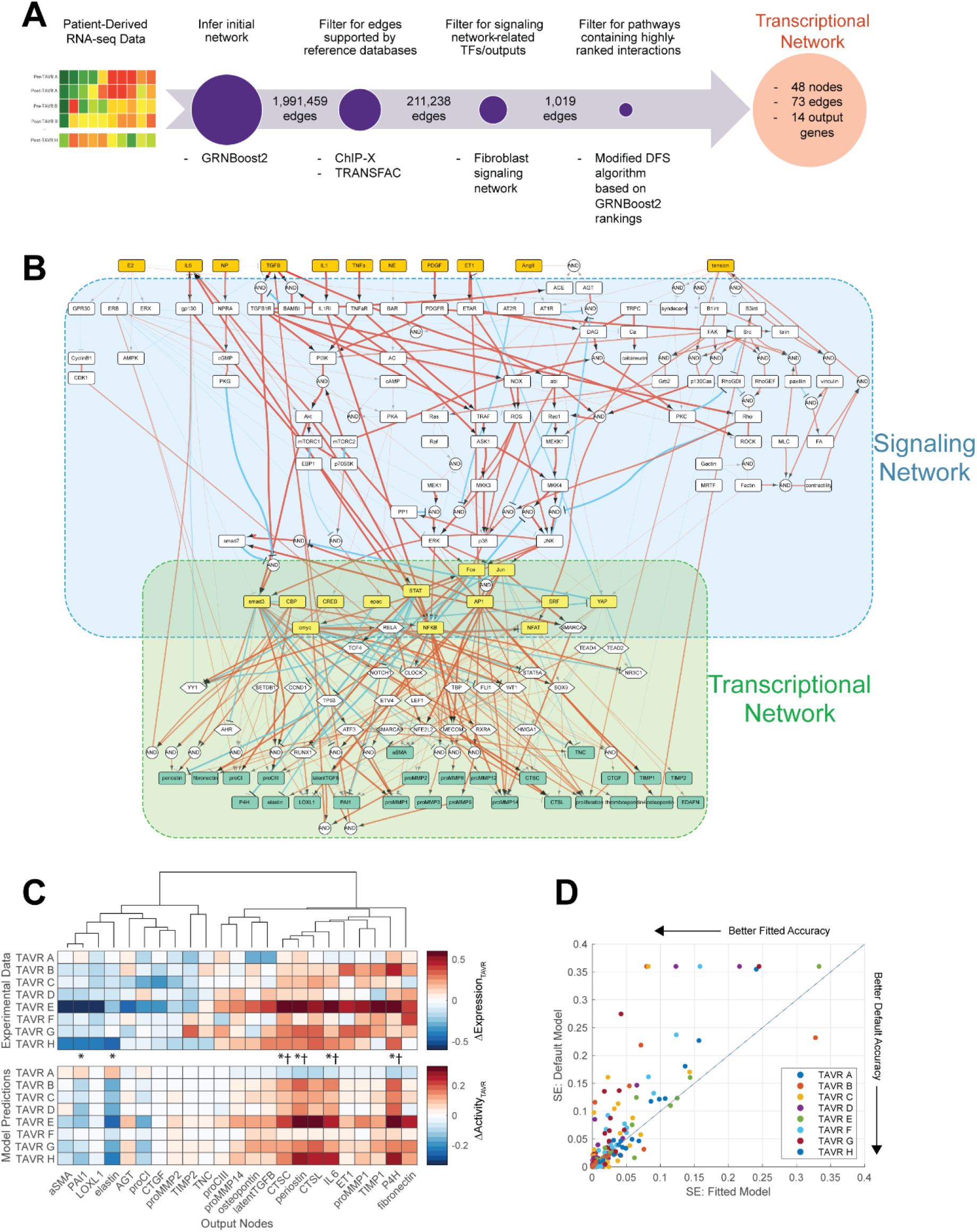
Composite myofibroblast regulatory network accurately predicts fibrosis-related protein expression based on patient-specific biochemical stimuli. (A) Schematic of gene regulatory network inference and model pruning scheme resulting in a targeted myofibroblast transcriptional network. (B) Topology of combined network consisting of a curated network of fibroblast signaling and an inferred transcriptional network derived from myofibroblast RNA-sequencing. Biochemical and biomechanical input nodes (orange boxes), signaling-activated TFs (yellow boxes), and fibroblast-secreted output nodes (green boxes) are connected by directed edges representing activating and inhibiting reactions (red and blue lines respectively, 334 total) with AND logic indicated by circular nodes. Line widths represent average reaction weight parameters estimated via genetic algorithm. (C) Comparison of experimentally measured changes in output gene expression between pre- and post-TAVR conditions (ΔExpression_TAVR_) with model predicted changes in output protein activity between pre- and post-TAVR conditions (ΔActivity_TAVR_). Individual comparisons made for each patient (boxes) represent mean levels across an ensemble of estimated parameter sets, and statistical comparisons of activity levels between pre- and post-TAVR groups were conducted via two-tailed Student’s t-tests. * p<0.05, † p<0.05 for model predictions and experimental values. (D) Comparison of squared errors (SE) of predicted patient ΔActivity_TAVR_ levels between fitted model (x-axis) and a default model (y-axis). All SE values were calculated against experimental ΔExpression_TAVR_ levels.

### Composite Myofibroblast Network Accurately Predicts Gene Expression

Myofibroblast activation and expression of ECM components during AVS relies not only on regulatory mechanisms at the transcriptional level, but also on changes in intracellular signaling that often involves crosstalk between multiple pathways. To provide a predictive solution that accounts for both levels of regulation simultaneously, we integrated the inferred transcriptional network above with our published network of fibroblast signaling, incorporating signaling reactions and TF-target interactions into a single composite network for modeling via a system of ordinary differential equations (Figure 1B). This composite network, consisting of 151 nodes and 334 edges in total, predicted the expression of 27 fibrosis-related proteins (output nodes) based on 11 canonical biochemical and mechanical stimuli (input nodes), providing a detailed footprint of myofibroblast responses to local environmental cues. This network was implemented as a system of logic-based ordinary differential equations (ODEs) as previously described^28^, in which normalized node activity levels are approximated via Hill equations using normalized node and reaction parameters (see Materials and Methods for full description).

We then contextualized this network to patient conditions before and after TAVR and maximized predictive capabilities of myofibroblast behavior during AVS progression. We implemented a genetic algorithm to estimate the relative weight parameters of all network reactions based on a published multi-omic dataset consisting of proteomic profiling of pre- and post-TAVR patient sera and RNA-sequencing of myofibroblasts cultured with patient-specific serum^12^. An ensemble of parameter sets was estimated using serum proteomic and myofibroblast transcriptomic datasets as analogs for model inputs and outputs respectively, and the resulting fitted network emphasized downstream signaling pathways from transforming growth factor-β (TGFβ), endothelin-1 (ET1), tumor necrosis factor-α (TNFα), and interleukins 1 and 6 (IL1 and IL6) as well as signal transducer and activator of transcription (STAT)- and nuclear factor kappa B (NFKB)-mediated transcriptional regulation (Figure 1B).

We compared model predictions of myofibroblast-expressed proteins to measured gene expression levels by simulating myofibroblast responses to each patient’s pre- or post-TAVR serum conditions and measuring changes in output activity between pre- and post-TAVR conditions (ΔActivity_TAVR_). Trends in model predictions matched changes in gene expression experimentally observed in pre- and post-TAVR serum-treated myofibroblasts (ΔExpression_TAVR_), with significant decreases in plasminogen activator inhibitor-1 (PAI1) and elastin activity along with significant increases in CTSC, CTSL, P4H, periostin, and IL6 activity mirroring significant changes in respective gene expression before and after TAVR (Figure 1C). Across the ensemble of estimated parameter sets, the average mean squared error (MSE) between ΔActivity_TAVR_ and ΔExpression_TAVR_ levels for each patient and model output was 0.0329 ± 0.0531, reaching a similar level as other fitted mechanistic and logic-based networks^29,30^.

We additionally assessed the accuracy of our parameter estimation strategy against a default model without fitted reaction weight parameters (i.e. all reactions weight parameters set to 1) by comparing ΔActivity_TAVR_ levels for each model against the previous ΔExpression_TAVR_ levels. We found that parameter estimation improved predictive accuracy for individual outputs with respect to each patient serum, as fitted model ΔActivity_TAVR_ levels for individual patients and outputs had lower squared errors against respective ΔExpression_TAVR_ levels compared to the default model (Figure 1D). Moreover, comparisons of significantly altered output levels between pre- and post-TAVR conditions as predicted by the two models and as measured in myofibroblasts suggests a sizeable improvement in predictive accuracy. We found that while the default model had a classification accuracy of 43.5% in predicting truly significant and non-significant changes in output activity compared to gene expression data, the fitted model improved the accuracy to 69.6%, thus predicting a higher proportion of truly significant and non-significant changes in output levels than the default, un-fitted model.

### Model-Predicted ECM Expression Correlates with Clinical Measures of AVS Severity

The relationship between fibrosis and reduced cardiac function during AVS is well-known, but linking individual secreted proteins to changes in valve and heart function remains challenging because scarring involves a variety of matrix, protease, matricellular, and crosslinking proteins for overall changes in tissue architecture. Our myofibroblast network can account for many of the species involved in scar formation, so we hypothesized that comparing model-predicted trends in myofibroblast protein expression to patient-matched clinical measurements could identify strongly correlated proteins among a diverse protein expression profile. We performed a systematic correlational analysis between activity levels of 23 outputs predicted for each pre-TAVR patient serum and 13 clinical features measured prior to surgery, utilizing serum levels of the patient cohort used for model training as well as levels for four additional patients as a validation set (Figure 2A). Several model-predicted outputs demonstrated significant positive correlations with left ventricle (LV) volume measurements (e.g. LV internal diameter at end diastole, LV internal diameter at end systole, and stroke volume index), including protease inhibitors TIMPs 1 and 2, matricellular proteins osteopontin and tenascin C, and autocrine feedback signals latent TGFβ, CTGF, ET1, and angiotensinogen. These associations suggest that high levels of feedback signaling and matricellular protein expression persist for patients with worse valve function and greater severity of AVS, although the underlying mechanisms remain unclear.

**Figure 2.**
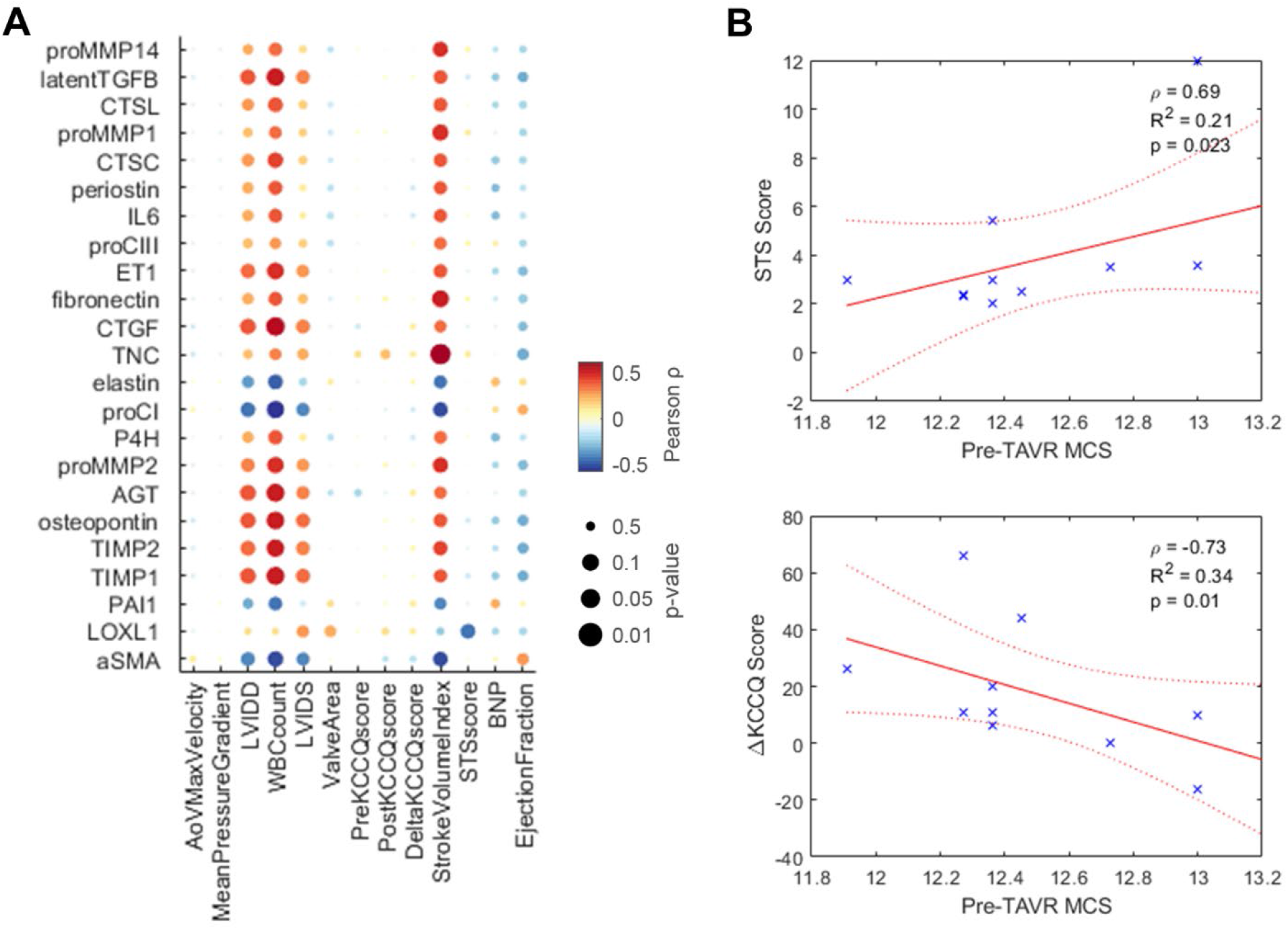
Model-predicted protein expression correlates with clinical measures of disease severity. (A) Correlation analysis of pre-TAVR output node activity levels (rows) with clinical features measured before and after TAVR surgery (columns). All dots represent Pearson correlation coefficients and respective p-values. (B) Notable correlations between aggregated matrix content score (MCS) and clinical features. Spearman correlation coefficients (ρ) and p-values were used to determine statistical significance.

We also aggregated individual output predictions into a single matrix content score (MCS) via a rank-based procedure as an analog for net matrix accumulation for each patient (see Materials and Methods for full description). Notable correlations include pre-TAVR MCS values with pre-TAVR Society of Thoracic Surgeons risk scores (STS score), and pre- to post-TAVR changes in Kansas City Cardiomyopathy Questionnaire score (ΔKCCQ score), which show that simulated ECM production is predictive of declines in ventricular function, overall disease severity, and negative post-surgical outcomes (Figure 2B).

### Network Perturbation Analysis Reveals Influential Pathways Mediating Fibrosis during AVS

A key barrier to understanding myofibroblast activation during AVS is a lack of insight into the relative contributions of individual signaling molecules across the full, multi-pathway network. Thus, we performed comprehensive knockdown simulations of individual nodes for all individual patient serum levels, measuring changes in network activity with each knockdown (ΔActivity_KD_) to assess network-wide effects in a patient-specific manner. We calculated a *knockdown influence* level for each node as a metric for the total effects that one node has on all other nodes, and we calculated a *knockdown sensitivity* level for each node as a metric for the total change in activity for one node across the knockdown of all other nodes (see Materials and Methods for full description). Selection of the top 10 scoring nodes in both metrics for each pre- and post-TAVR serum condition showed similar trends in node sensitivity and influence across patients, with a total of 19 nodes ranking in the top 10 knockdown influence and 17 nodes ranking in the top 10 knockdown sensitivity for at least one of the eight patients (Figure 3A-B, see Figure S2A for network-wide sensitivity analysis).

**Figure 3.**
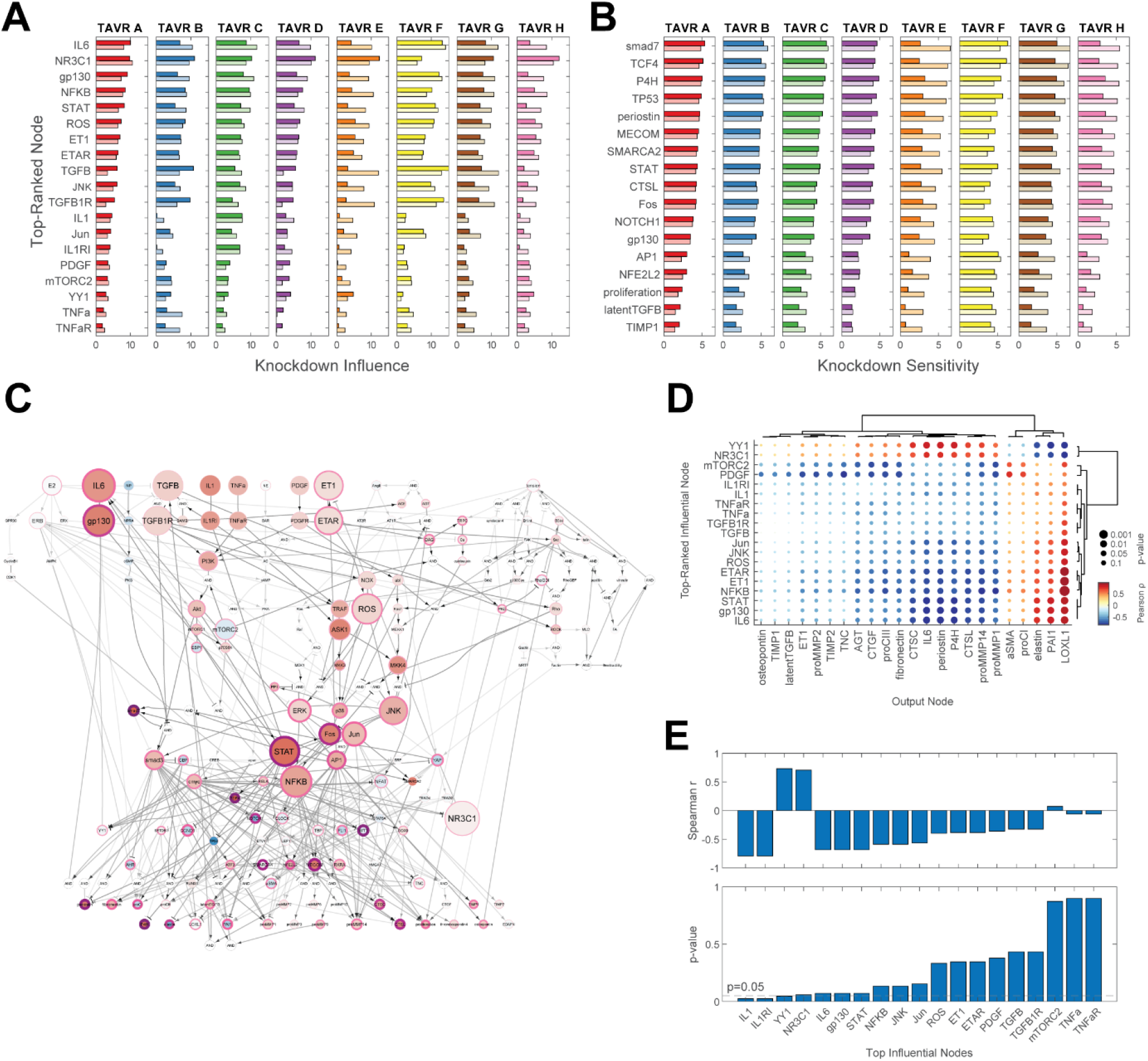
Network perturbation analysis reveals influential pathways and differences in myofibroblast responses across patient conditions. (A-B) Comparison of top-ranked influential and sensitive nodes between patients and pre-/post-TAVR conditions. Nodes shown reflect all unique nodes ranked in the top 10 based on knockdown influence (A) or knockdown sensitivity (B). Dark bars represent knockdown influence/sensitivity levels for each pre-TAVR patient condition, and light bars represent levels for each post-TAVR patient condition. (C) Topological representation of influential and sensitive pathways within the myofibroblast network. Node sizes represent average knockdown influence levels across all patient and pre-/post-TAVR conditions, node colors represent average ΔActivity_TAVR_ levels across all patients, and node border sizes/colors represent average knockdown sensitivity levels across all patient and pre-/post-TAVR conditions. (D) Correlational analysis between pre-TAVR knockdown influence levels for top-ranked nodes and ΔActivity_TAVR_ levels for model outputs. All dots represent Pearson correlation coefficients and respective p-values. (E) Correlational analysis between knockdown influence levels for top-ranked nodes and predicted ΔMCS levels between pre- and post-TAVR conditions.

Across all patient and serum conditions, several intracellular signaling pathways were consistently influential with respect to network-wide activity. Among the top influential pathways represented were IL6 signaling via gp130/STAT activation, canonical and noncanonical TGFβ signaling via smad3 and ROS, extracellular signal-regulated kinase (ERK), and JNK activation, and ET1 signaling via ROS/ERK/JNK activation along with minor influence via IL1, TNFα, and PDGF signaling (Figure 3C). Nodes downstream from these pathways consistently scored highly in knockdown sensitivity, including targets of IL6 signaling such as smad7, TCF4, TP53, and NOTCH1 as well as targets of TGFβ/ET1 signaling such as AP1, NFKB, cmyc, and MECOM. These influential and sensitive species coincide with sizeable changes in activity between pre- and post-TAVR conditions, as average ΔActivity_TAVR_ levels across all patients indicate upregulation of IL6 pathway activation after TAVR and large increases or decreases in downstream activity including connected TFs and target output nodes (Figure 3C).

We next compared knockdown influence and sensitivity levels for top-ranked nodes across patient-specific conditions to assess differences both between individual patients and between pre-/post-TAVR conditions. We found that influential nodes representing IL1 and TGFβ signaling pathways fluctuated between the patient cohort, as select patient conditions showed large network-wide changes in activity compared to others (Figure 3A), and calculated coefficients of variation across patient conditions were largest for IL1 and TGFβ-associated nodes out of all top-ranked nodes (Figure S2B). While patient-specific differences in knockdown sensitivity levels were not as pronounced and coefficients of variation were generally lower (Figure S2C), overall knockdown sensitivity levels did vary by patient with some patients showing little sensitivity across all top-ranked nodes compared to others (Figure 3B). We also observed differences in influence and sensitivity from pre-TAVR to post-TAVR conditions across top-ranked nodes. Knockdown influence levels for IL6 signaling nodes increased for post-TAVR conditions compared to pre-TAVR conditions for 7 of 8 patients, which was consistent with increases in IL6-related activity during post-TAVR simulations.

As an additional exploratory analysis of influential pathway effects on myofibroblast behavior, we correlated pre-TAVR knockdown influence levels of top-ranked nodes with ΔActivity_TAVR_ levels for all fibrosis-related outputs to discover relationships between pre-TAVR conditions and changes in cell behavior with surgical intervention. A total of 82 out of 437 correlations between pre-TAVR influence and output ΔActivity_TAVR_ levels were significant including 27 positive correlations and 55 negative correlations (Figure 3D). Within these pairings, influence levels of IL6-associated nodes and ET1-associated nodes significantly correlated with nodes that ranked highly in knockdown sensitivity (Figure 3D). Positive correlations with elastin, PAI1, and LOXL1 as well as negative correlations with multiple matrix proteases, periostin, and P4H suggest strong connections between pre-TAVR pathway knockdown and changes in protease, protease inhibitor, and collagen crosslinking protein expression over other ECM-related proteins. We additionally correlated pre-TAVR influence levels with changes in MCS values from pre-TAVR to post-TAVR (ΔMCS) as a metric for overall changes in myofibroblast-mediated matrix turnover, and we found that knockdown influence of the IL1- and ET1-associated nodes positively correlated with changes in matrix content for patients (Figure 3E), suggesting that these pathways may contribute to overall changes in myofibroblast protein secretion within the larger network.

### Targeted Virtual Drug Screen Predicts Stratified Patient Responses to Therapies

Our perturbation analyses above suggested that while some signaling pathways tend to mediate myofibroblast protein expression over others, responses of individual patients to pathway perturbations can vary widely with some patients experiencing little to no benefit for a given therapy. This variability is a common challenge for cardiovascular therapy development, so we investigated whether our myofibroblast network could discern patient-specific responses to drugs by predicting personalized changes in fibrosis-related outputs. Inhibition of each node was performed for each patient’s pre-TAVR conditions in a dose-response manner by limiting of the maximum activation of each node, and average changes in activity of output nodes (ΔActivity_inhib_) were measured for each patient relative to unperturbed levels. This knockdown analysis identified the 14 most influential nodes (Figure 4A rows) and the 15 most sensitive outputs (Figure 4A columns) on average, but also demonstrated wide distributions of patient responses to each perturbation. As an example, inhibiting endothelin-1 receptor (ETAR) produced either minimal, moderate, or substantial reductions in OPN production as well as minimal, moderate, or substantial increases in PAI1 production depending upon the particular patient serum context (Figure 4B). In addition, different patient backgrounds exhibited different response sensitivity depending on the protein output of interest.

**Figure 4.**
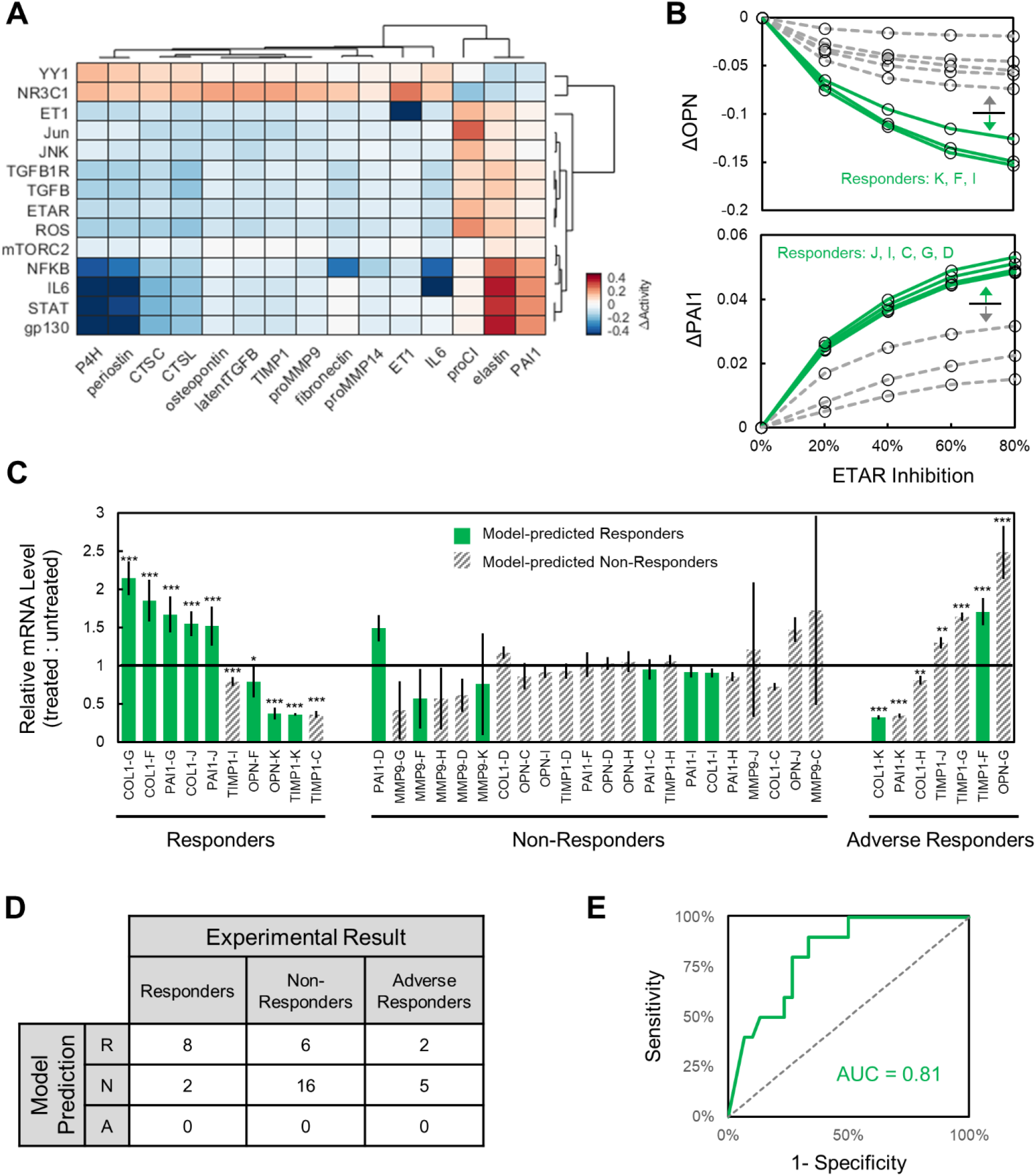
Virtual drug screen reveals stratified patient responses to inhibitor treatments. (A) Comparison of ΔActivity_inhib_ levels for fibrosis-related model outputs upon knockdown of top-ranked influential nodes. Values represent average ΔActivity_inhib_ levels across all patients for pre-TAVR conditions after 80% inhibition of each node relative to respective un-perturbed conditions, and rows/columns were chosen for knockdowns that demonstrated an average 5% change in activity for at least 5 outputs and vice-versa. (B) Simulated changes in OPN and PAI1 output activity for each patient with 0 to 80% inhibition of ETAR. Patients were designated as ‘Responder’ or ‘Non-Responder’ based on threshold classification of delta levels at 80% inhibition. (C) Relative in vitro gene expression levels of OPN, PAI1, COL1, MMP9, and TIMP1 across valve interstitial cell populations treated with patient-specific sera and ETAR inhibition with bosentan. Gene expression was measured via qPCR relative to RPL30 for cells dosed with individual patient sera for 24h followed by 10 μM bosentan for 24h, and statistical significance is shown as * p<0.05, ** p<0.01, ***p<0.001. (D) Confusion matrix comparing model predictions of patient responder/non-responder status to that determined via qPCR results. (E) Receiver-Operating-Characteristic curve comparing demonstrated predictive accuracy of model predictions vs. in vitro patient-specific response status.

To experimentally validate the predictive accuracy of our model to capture patient-specific drug responses, we subjected porcine VIC populations to the ETAR inhibitor bosentan across 8 patient-specific serum culture conditions (see Materials and Methods for full description). Briefly, we cultured cells on poly(ethylene glycol) hydrogels that served as precision biomaterials^31^ and then exposed the cell-laden hydrogels to individual patient sera with or without 10 μM bosentan. After 24h of treatment, we collected mRNA and measured OPN, PAI1, COL1, MMP9, and TIMP1 expression levels using RT-PCR (Figure 4C). Patients were first classified as ‘responders’ or ‘non-responders’ for each protein output based on computationally predicted effects of ETAR 80% knockdown and a simple over-under threshold. Computational model-based classifications across patient backgrounds and genes were compared to experimental results for patient backgrounds and genes by designating each experimental result as a responder (i.e., statistically significant effect of bosentan in the same direction as the model prediction), an adverse responder (i.e., statistically significant effect of bosentan in the opposite direction as the model prediction), or a non-responder (i.e., statistically insignificant effect of bosentan in any direction). The model-based stratification showed substantial predictive power with 8/10 experimental responders correctly identified in the computational responder group while only 6/16 non-responders and only 2/7 adverse responders were incorrectly classified by the model as responders (Figure 4D-E). Varying the model stratification threshold produced a full receiver-operating-characteristic curve with an area-under-the-curve of 0.81.

## Discussion

Preventing valve fibrosis during AVS is crucial for long-term patient outcomes even after valve replacement interventions. Although previous studies have shown that alterations in serum protein levels after valve replacement can deactivate myofibroblasts^12,13^, the underlying mechanisms of this deactivation remain unclear due in part to the complexities of intracellular regulatory mechanisms. Using coupled serum proteomic and myofibroblast transcriptomic datasets for patients undergoing TAVR, we developed a strategy to integrate this patient-specific data into a computational model of myofibroblast signaling and transcriptional regulation to predict fibrosis-related protein expression in response to environmental signals. We found that this strategy improved the accuracy of patient-specific predictions over simulations based on model topology alone and resulted in a similar degree of accuracy against experimental gene expression data compared to other fitted network models^29,30^. Correlational analyses demonstrated strong trends between both individual matrix proteins predictions and aggregated matrix content scores with clinical measures of disease severity and patient quality-of-life. Network-wide perturbation analyses identified influential signaling pathways and transcriptional regulators of network activity and output expression such as the IL6-gp130-STAT, TGFβ-ROS-JNK, and ET1-ROS-ERK/JNK signaling axes. We additionally demonstrated the capacity of the model to detect stratified patient responses to targeted inhibition in a virtual drug screen, showing that myofibroblasts respond substantially to simulated inhibitors for select patient conditions over others. Our results suggest that this strategy can accelerate the identification of anti-fibrotic therapies to slow AVS progression by accounting for individual patient conditions and generating insight into possible mechanisms governing cell behavior within a complex signaling context.

### Identification of Transcriptional Regulators in Myofibroblast-Mediated Fibrosis

While intracellular signaling processes remain a crucial step in transducing environmental cues into changes in cell protein expression and phenotype, regulation at a transcriptional level adds an additional layer of complexity and can confound understanding of cell behavior. Our construction and simulation of a myofibroblast transcriptional network derived from cell RNA-sequencing data suggest that this layer of regulation plays a highly influential role in myofibroblast behavior during AVS progression. Transcriptional regulators identified here via gene regulatory network inference have also been shown to mediate VIC and cardiac fibroblast behavior in other cardiovascular pathologies. The transcription factor SOX9 has been identified as a positive regulator of cardiac fibrosis and heart failure development in models of ischemia-reperfusion injury and myocardial infarction (MI), with ischemia causing increased SOX9 gene expression and spatial correlation with collagen gene expression and scar formation measured via histology^22^. Fibroblast-specific deletion of SOX9 in an MI mouse model has also been shown to attenuate increases in αSMA expression, cell proliferation, scar formation, and functional measures. These data demonstrate a strong connection between this factor and fibrotic processes.

NOTCH1 and ATF3 have also been implicated in cardiac fibrosis and myofibroblast activation, as overexpression of the NOTCH1 intracellular domain in rat cardiac fibroblasts antagonized TGFβ-induced smad3 activation and *in vivo* overexpression after MI reduced fibrotic area in rats^27^. Further, cardiac fibroblast-specific overexpression of ATF3 attenuated scar formation and reduced the LV internal diameter in end-systole and ejection fraction after MI^26^. It should be noted that several regulators identified by our approach serve influential roles in osteogenic differentiation such as NOTCH1^32–34^, and although the scope of our model does not directly reflect osteogenic phenotypes for valvular cells, this similarity in regulatory species suggests that pathways mediating myofibroblast and osteoblast phenotypes are not mutually exclusive. Our identification of these transcriptional intermediates and connecting interactions suggests that the regulation of myofibroblast behavior involves numerous pathways simultaneously, and that computational approaches are advantageous for discerning mechanisms essential to cell function within a dense regulatory landscape.

### Influential Signaling Axes and Regulatory Hubs Mediate Overall Myofibroblast Behavior

Myofibroblasts respond to a variety of environmental stimuli including inflammatory cytokines, growth factors, matrix stiffness, and hormonal peptides, all of which have been shown to alter cell behavior individually^7,35^. Within the context of AVS progression, however, the relative contribution of each environmental signal and their respective signaling pathways remains unclear. We performed a network-wide perturbation analysis to identify globally influential pathways with cells cultured in the presence of pre-TAVR and post-TAVR serum, and our simulations predicted that the transduction of IL6, TGFβ, and ET1 primarily influence activation levels across the network. IL6 signaling and downstream activation of STAT has been previously linked to cardiac fibrosis across several cardiovascular pathologies with effects across multiple pathways^36^.

Dziemidowicz and colleagues found that IL6^−/−^ mice treated with isoproterenol developed interstitial ventricular fibrosis and increased phosphorylated levels of multiple signaling intermediates including ERK, Raf, and p38 compared to wild-type mice^37^. In a similar finding, Hilfiker-Kleiner and colleagues found decreases in ERK and Akt phosphorylation for mice carrying a point mutation in gp130 and subjected to MI compared to wild-type mice. Recent evidence of IL6 as a promoter of VIC osteogenic differentiation via the increased expression of RUNX2 and osteopontin suggest that this pathway may serve multiple phenotypic roles^13^, and future targeted investigations of this behavior may explain possible context-dependent roles of IL6 signaling during AVS progression.

Noncanonical TGFβ signaling via NADPH oxidases (NOX) and ROS has also been previously revealed as an effective regulator of fibrosis in myocardial and valvular tissue, supporting the high influence that we found from the TGFβ-ROS-JNK axis in our computational network. Inhibition of NOX4 in cardiac fibroblasts has been shown to reduce TGFβ-stimulated intracellular superoxide production, which in turn mediates phosphorylation of smad2/3 and expression of αSMA, CTGF and fibronectin, suggesting a largely pro-fibrotic role of this pathway^38^. The NOX-ROS signaling axis has also been shown to mediate changes across multiple pathways to alter fibrosis in mouse models of hypertension^39^, and decreases in NOX4 expression and ROS generation following genetic knockout of SOD1 in mice mitral valves has been associated with changes in αSMA, CTGF, and MMP expression in mitral VICs^40^, suggesting a similar systemic effect of this noncanonical mechanism on valvular myofibroblast behavior.

Our global analysis additionally suggests that select TFs may act as regulatory hubs for expression of ECM-related proteins and influence fibrotic behavior over others. We found that STAT, NFKB, and Jun influence large changes in network-wide activity, all of which have been well-studied for their roles in pathological scar formation. In particular, we found STAT to be a primary influencer of myofibroblast behavior via the expression of matricellular proteins, cathepsin and MMP proteases, and crosslinking enzymes, and previous experimental studies have confirmed this central role of STAT in cardiac fibrosis and systemic sclerosis. Using IL-11 as a positive regulator of STAT3 after MI in mice, Obana and colleagues demonstrated that STAT3 activation reduced LV scar formation and improved infarct wall thicknesses compared to un-treated MI, demonstrating a primarily anti-fibrotic effect during ischemic injury^41^. However, disruption of STAT3 phosphorylation via inhibition of EphrinB2 in a post-MI mouse model reduced αSMA protein expression, collagen I expression, and cell proliferation^42^, and inhibition of STAT3 during bleomycin-induced skin fibrosis reduced gene expression of multiple ECM proteins^43^, suggesting that STAT may exert both pro- and anti-fibrotic effects. In agreement with experimental evidence of IL6 signaling during AVS, the activation of STAT3 was also associated with osteogenic de-differentiation in VICs via downregulation of RUNX2 and BMP-2^44^, reinforcing the need to further investigate this intersection between fibrotic and osteogenic phenotypes.

### Systems Models Relate Cell Phenotypes to Disease Severity and Predict Patient Responses

Our model-centered approach provides unique advantages for relating cellular-level behavior to clinical manifestations of AVS through the prediction of protein expression based on patient-specific conditions. Our findings indicate that computational predictions of cell behavior correspond with multiple measures of AVS severity including LV size and STS scores. Previous serum biomarker studies in AVS cohorts have shown individual biomarkers to be predictive of patient outcomes such as N-terminal pro B-type natriuretic peptide (NT-proBNP) and C-reactive protein^45,46^, and while useful for stratifying patient risk and informing surgical intervention, these biomarkers alone may not connect alterations in cellular function with overall disease progression. Our patient-specific predictions and correlation with pre-TAVR clinical measurements suggest that matricellular protein expression, autocrine feedback expression, and aggregation of all output proteins into a single ECM-related metric offer a predictive link between myofibroblast-mediated scar formation and reduced heart function. Virtual drug screens for individual patient conditions further demonstrated the ability of this model-based approach to discern responders versus non-responders toward possible therapies. Translating this approach into a clinical setting may improve overall responses to therapeutic intervention before and after surgery via targeted drug and/or dose selection. While clinical validation of these trends is essential for further development and application of this approach, the framework developed here provides opportunities to further investigate fibrotic mechanisms-of-action for targeted patient cohorts or for other possible pathological scenarios.

### Study Limitations

Our study was primarily limited by the size of patient datasets used for model training and simulated scenarios. While this multi-omic, patient-matched design formed an essential aspect of our model development strategy through the robust measurement of serum proteins and cell-specific gene expression, a lack of cohort size and diversity limited our predictive capabilities in terms of both model training and predicted trends in myofibroblast behavior. Additionally, the insufficient availability of select biochemical stimuli (e.g. estrogen), tissue biomechanical properties, and cell-secreted outputs limited our predictions related to these stimuli and associated regulatory pathways. In particular, the signaling network model was curated with substantial mechano-transduction pathways enabling the incorporation of local biomechanical signals such as tissue deformations and stiffness as an additional input which was not leveraged in these current simulations.

Because both fibrosis and calcification processes play key roles in the development of AVS, we acknowledge that our model of myofibroblast protein expression alone accounts for only one component mediating overall disease progression. Future adaptation of this modeling strategy towards osteogenic differentiation may provide additional insight into dominating mechanisms responsible for calcification and complement our current model of fibrosis-related gene expression for an overall picture of cell behavior. Because several influential pathways identified here also correspond with the expression of calcification-related proteins, implementing these same pathways along with those known to mediate severe AVS may be valuable for investigating differences in fibrotic versus osteogenic differentiation.

### Conclusion

Our development of a fibroblast regulatory network and analysis of patient-specific changes in myofibroblast protein expression demonstrated the translational possibilities of model-based approaches in identifying key mechanisms governing cell behavior within a larger signaling context. Predicted intracellular species and pathways influencing fibrosis-related gene expression provide targeted hypotheses for future clinical validation. Our findings of strong correlations between model-predicted protein expression and clinical measures of disease severity suggest that simulations of biological networks can be advantageous for relating cellular phenotypes to clinical manifestations of disease, and future exploration of the predictive capabilities of such models could prove useful for clinical translation. Our virtual drug screen of individual patient responses to pathway-inhibiting drugs further demonstrated the ability of this model-based approach to stratify patient cohorts based on individual serum biomarker levels. This strategy provides a framework for predicting cell behavior within a complex signaling context and enables further investigation into pathways mediating valvular fibrosis before and after surgical intervention.

## Acknowledgments

B.A.A. acknowledges funding from the National Institutes of Health (R00HL148542) and the Burroughs Wellcome Fund Postdoctoral Enrichment Program, K.S.A. acknowledges funding from the NIH (R01HL132353 and R01HL142935), and W.J.R. acknowledges funding from the NIH (R01HL144927 and P20GM121342).

## Materials and Methods

### Data and Code Availability

The code generated during this study is available at GitHub (https://github.com/jdroger/Fibroblast_Gene_Regulatory_Network). The published article includes all models generated or analyzed during this study.

### Fibroblast Transcriptional Network Construction

#### Gene Regulatory Network Inference

We employed a previously published method of gene regulatory network inference to derive a network of transcription factor (TF)-target gene interactions based on a transcriptomic dataset of VIC populations after stimulation with patient sera. Bulk RNA-sequencing datasets recently published by our collaborators were used as a training set for network inference, and this dataset represents the gene expression of VIC populations stimulated with serum derived from a small cohort of patients undergoing a TAVR procedure^12^. Briefly, patient blood samples were collected at the time of surgery and at the 1-month follow up visit (n = 4 female patient pairs and n = 4 male patient pairs, 16 samples total). The resulting serum samples were used to treat porcine VIC cultures seeded on soft PEG hydrogels (Young’s modulus of 5.8 kPa) for 48 h prior to RNA-sequencing, and counts per million (CPM) for gene expression were used for subsequent network inference and model fitting methods.

We utilized the GRNBoost2 machine learning algorithm to infer a network of TF-target interactions from the transcriptomic dataset above. This regression-based method is based on the GENIE3 tree-based algorithm for predicting regulatory links between input genes and target genes via the construction of decision tree ensembles^47^. Each ensemble of decision trees, which predicts the expression of a given target gene from the expression of all input genes, is used to determine the relative “*importance*” of each input gene in predicting the expression of the specified target gene. Decision tree ensembles are built for all genes across the transcriptome, and input-target gene links are aggregated to form a composite network of ranked interactions. The GENIE3 algorithm has been shown to out-perform other methods in inferring gene regulatory networks as part of the DREAM4 *In Silico Multifactorial* challenge^47^, and it provides several advantages over other common inference algorithms: (1) inference can be performed with minimal assumptions of network topology, (2) directed interactions (i.e. gene A activates gene B) can be inferred compared to correlation- and probability-based methods, and (3) non-linear or combinatorial regulation can be derived compared to other regression-based methods^48^. The GRNBoost2 implementation optimizes this approach using stochastic gradient boosting, which grows decision tree ensembles on a subset of observations and estimates the loss function on the remaining observations with each iteration. The algorithm implements an early stopping criterion if the loss function does not improve above a set threshold, thereby preventing unnecessary iterations of each decision tree and reducing overall computational time^49^. The Arboreto library for python was used to apply this algorithm to the RNA-sequencing data described above, and the average computational time for network inference using a 4-core computer was approximately 10 min.

#### Network Pruning

Upon inference of the initial network, a 3-step workflow was applied to filter the network for TF-target interactions that satisfy 3 requirements: (1) interactions must be supported by experimental evidence, (2) interactions must relate to either literature-supported TFs (primary TFs) or fibrosis-related target genes in myofibroblasts, and (3) resulting pathways connecting primary TFs and target genes must have relatively strong links across all individual edges as determined by interaction ranks assigned by the GRNBoost2 algorithm (Figure 1A). The first step was performed by comparing inferred TF-target interactions with databases of known TF-target interactions aggregated from chromatin immunoprecipitation (ChIP) studies. Curated lists of known TF-target interactions from the ChIP-X Enrichment Analysis (CHEA) and TRANSFAC databases were downloaded using the Harmonizome web interface^50^, both of which were chosen to maximize coverage of TFs related to myofibroblast activation. Upon filtering the initial gene regulatory network for interactions contained in either database, the resulting network was filtered further for interactions containing TFs or fibrosis-related target genes in myofibroblasts. Lists of TFs and target genes were derived using our curated network model of fibroblast signaling (see Chapter 4), which contains 11 primary TFs and 20 target genes coding for ECM-related proteins. For TFs that consist of several subunits (e.g. activator protein 1 [AP1] complex), both constituent genes were included for filtering. An additional 8 target genes that were differentially expressed by patients between pre- and post-TAVR sera were also considered: CTSC, CTSL, COL4A5, LOXL1, P4HA1, P4HA3, LAMA4, and ELN. Proteins coded by these genes have been shown to alter matrix degradation, collagen processing, and alter material properties of cardiac tissue, and significant differences in expression within the RNA-sequencing dataset provide a rationale for exploring possible regulatory pathways affecting gene expression. After list construction, the database-filtered network above was filtered again for interactions containing genes in either list. After filtering, interactions contained only TF-TF interactions or TF-target interactions in which intermediate TFs not included in the primary TF list above regulate target gene expression (i.e. primary TF A activates secondary TF B, which activates target gene C).

After the second stage of filtering above, the final network topology was derived by ensuring that all resulting pathways between primary TFs and target genes contained interactions that ranked highly among possible regulatory links according to the GRNBoost2 algorithm. A modified depth-first-search algorithm was implemented to find all possible pathways between each primary TF and target gene and check whether each interaction within that pathway met this requirement using the “*importance*” score for the interaction output by GRNBoost2. Individual interactions were only allowed if the importance score of each edge was greater than either a threshold of 1 or the 75^th^ percentile of all interactions stemming from the same TF. This hybrid threshold was chosen to both limit interactions driven by noise, in which overall importance scores are low, and to prevent premature exclusion of related interactions when all possible regulatory links may be ranked low relative to the entire network. By implementing this method, all interactions mediating expression of target genes were ensured to meet a threshold of confidence relative to neighboring interactions such that one TF within a pathway is not predictive of its downstream target. All network filtering steps were performed in a python environment using the numpy^51^ and pandas^52^ packages.

### Composite Signaling/Transcriptional Network Implementation

#### Topological Integration

We combined the final transcriptional network with our previous fibroblast signaling network describing intracellular mechanotransduction and chemotransduction (see Chapter 4) to form a composite network capable of predicting fibrosis-related protein expression in response to mechanical and biochemical stimuli. New TFs (model nodes) and/or transcriptional reactions (edges) were added to the fibroblast signaling topology if they were not redundant to the original transcriptional reactions described by the signaling network.

#### Logic-Based Ordinary Differential Equation Approach

The final network was implemented as a system of logic-based ordinary differential equations in which activity levels of all nodes were modeled by Hill equations. Logical NOT, AND, and OR gates were used for complex signaling interactions by applying the respective logical operations: 1 − *f*(*x*) for NOT gates, *f*(*x*) *f*(*y*) for AND gates, and *f*(*x*) + *f*(*y*) − *f*(*x*) *f*(*y*) for OR gates. The open-source Netflux package for MATLAB was used to build this system of differential equations^28^, and all simulations were conducted using MATLAB (Mathworks, Natwick MA). All visualizations of network topology were constructed using Cytoscape^53^.

### Network Parameter Estimation

#### Dimensionality Reduction

To improve model predictions of myofibroblast behavior within the context of AVS before and after TAVR, we implemented a model fitting procedure to optimize the reaction weight parameters (w) of all reactions within the composite network. From the initial set of parameters (334 total), k-means clustering was conducted based on a global sensitivity analysis to group reaction weights based on changes in network-wide activity with knockdown. This method has been utilized in previous logic-based ordinary differential equation models to identify modules with similar functional behavior^17^ and provides an advantageous method for reducing model dimensionality based on biological function. Reactions were clustered via the *kmeans* MATLAB function using k = 11, which was determined to produce the highest degree of separation between clusters. Reactions in which the product contained a fibrosis-related output gene were excluded from clustering to allow for additional degrees of freedom to predict the expression of individual output proteins, and k-means clustering was repeated 1000 times for the remaining reactions using randomly chosen starting points. Clusters of reactions occurring most frequently were assigned to final clusters and shared a weight parameter during fitting, thus reducing the dimensionality of the final parameter set for optimization (129 parameters total).

#### Multi-Omic Data Normalization

A genetic algorithm was used to fit the reduced parameter set above to normalized model input and output concentrations extracted from patient-specific proteomic and transcriptomic datasets recently published by our collaborators^12^. In addition to the VIC RNA-sequencing data described above, relative concentrations of 1193 proteins from the same patient serum samples were measured via DNA aptamer array, allowing for the direct relation of changes in serum proteins before and after TAVR to changes in myofibroblast activation. Using serum protein levels for all patients as input concentrations and VIC gene expression levels as output concentrations, data for each input and output node were transformed to a normalized scale useable by the model. All input levels were normalized between activity levels of 0.1 and 0.6 (representing 10% activation and 60% activation respectively), matching previous studies transforming biochemical cytokine and growth factor levels measured in myocardial tissue post-MI to normalized values while maximizing network dynamic range^54^. All output levels were normalized between activity levels of 0.1 and 0.7, keeping the same basal values as input levels while expanding the dynamic range of output expression beyond the default half-maximal effective concentration (EC50) to account for multi-input stimulation. All normalized input levels were implemented as initial reaction weights within the model as representative rates of generation of each species, and all normalized output levels were implemented as steady-state concentrations of each species. Because not all input/output nodes were represented in the experimental datasets, unrepresented input reaction weights were included as parameters within the fitting parameter set, and unrepresented output activity levels were included for prediction but not used in the final objective function.

#### Global Parameter Estimation

Using normalized input/output sets for each patient, the *ga* MATLAB function was used for all parameter estimation. For each set of randomly generated parameter sets within the function, steady-state output levels for each patient were measured after 80 h given either pre-TAVR or post-TAVR levels for that patient. Changes in steady-state output levels were calculated for each patient from pre-TAVR levels to post-TAVR levels (ΔActivity_TAVR_), and the mean squared error (MSE) between ΔActivity_TAVR_ values and experimental changes in normalized gene expression between conditions (ΔExpression_TAVR_) for all patients and outputs were used as the objective function. A population of 500 was chosen to maximize intra-generational variation while limiting computational resources, and a maximum of 100 generations were used as minimal improvements in the objective function were found in subsequent generations (data not shown). Program defaults were used for all other algorithm hyperparameters, and default values for the fibroblast signaling model were used for all other model parameters including Hill coefficient (n), EC50, maximum node activation (Y_max_), and time constant (⊺) (see Chapter 4, Materials and Methods section for full description). Due to the stochastic nature of generating initial parameter sets for global optimization, the model fitting procedure was repeated 50 times using random initial parameter sets resulting in an ensemble of parameter sets. Predicted node activity levels for all subsequent simulations reflect the mean activity levels predicted across all estimated parameter sets.

### Network Perturbation Analysis

To identify influential signaling mechanisms across pre-TAVR and post-TAVR signaling contexts, a series of node knockdowns were simulated using normalized input levels from each patient serum sample. For each set of normalized input levels used during parameter estimation above, basal conditions (i.e. without any knockdown) were applied for 80 h, followed by knockdown of individual nodes using the Y_max_ parameter (Y_max,KD_ = 0.1*Y_max,basal_) for 240 h. Steady-state activity levels of all nodes were measured with each knockdown, and changes in node activity (ΔActivity_KD_) were calculated as the difference between node activity after knockdown and basal node activity. Knockdown sensitivity of each node was calculated as the sum of absolute ΔActivity_KD_ levels for the node across all knockdown simulations, and knockdown influence of each node was calculated as the sum of absolute ΔActivity_KD_ levels for all other nodes in the network upon knockdown.

### Patient stratification analysis

Targeted simulations assessing stratified model responses to drug targets with patient-specific conditions were conducted using a series of dose-response simulations. For each node, a series of knockdown simulations was performed under each patient-specific condition. Following a simulation of basal conditions using each patient’s normalized input levels for 80 h, the Y_max_ parameter for each node was lowered to either 0.8, 0.6, 0.4, or 0.2 times the Y_max_ under basal conditions for 240 h (corresponding with 20-80% node inhibition). Steady-state activity levels of all nodes were measured with each dose and compared to steady-state levels prior to dosing. Patients were then classified as ‘responder’ or ‘non-responder’ based on a simple over-under threshold, which was varied from 0 to the maximum delta for each protein output to generate receiver-operating-characteristic curves.

### In vitro validation experiments

#### PEG Hydrogel Fabrication

Poly(ethylene glycol) (PEG) hydrogels were made as previously described^12^. 25 mm glass coverslips were O_2_ plasma-treated and treated in a 15% vol/vol mercaptopropyltrimethoxysilane (MPTS, Sigma-Aldrich) and 5% vol/vol 2-butylamine (Sigma-Aldrich) solution in toluene (Sigma-Aldrich) for 2 hours to functionalize the glass with free thiols. Coverslips were rinsed with toluene, dried in a 80°C oven, and sterilized with 70% ethanol. Gel precursor solutions (4% wt/vol PEG-norbornene [Nb]) were prepared by mixing 8-arm 40 kDa PEG-Nb with 5 kDa PEG-dithiol crosslinker (JenKem) and 2 mM CRGDS cell adhesive peptide (Bachem) at a 0.99:1 thiol-to-ene ratio in phosphate buffered saline (PBS, Sigma-Aldrich). Lithium phenyl-2,4,6-trimethylbenzoylphosphinate (LAP, 1.7 mM) was added to the pre-cursor solution prior to photo-polymerization. The gel pre-cursor solution was sandwiched between a coverslip and a Sigmacote (Sigma) treated microscope glass slide (final gel thickness = 150 μm for 25 mm coverslips), UV-photopolymerized at 4 mW/cm^2^ for 3 minutes, sterilized in 5% isopropyl alcohol in PBS for 30 minutes, washed 3 times with PBS, and swelled overnight in VIC media at 37°C and 5% CO_2_.

#### Rheology

The PEG hydrogel precursor solution (4% wt/vol PEG-Nb formulation) was photopolymerized *in situ* on a HR3 rheometer (TA Instruments) and characterized using a parallel plate geometry (8 mm diameter). Storage (G’) and loss (G”) measurements were made using oscillatory shear rheology with an amplitude of 1% and frequency of 1 Hz. The shear modulus values (Figure S3) were converted to Young’s modulus (E) using the formula E = 2G’(1+ν) with a Poisson’s ratio of ν = 0.5, assuming G’ ⋙ G”.

#### VIC isolation and culture

Male or female porcine valvular interstitial cells (VICs) were harvested from porcine aortic valve leaflets as previously described^12,55^. Briefly, aortic valve leaflets were excised from fresh porcine hearts from 5-6-month-old young adult pigs (Hormel), rinsed in warmed Earle’s Balanced Salt Solution (EBSS, Sigma-Aldrich) supplemented with 50 U/mL penicillin (Thermo Fisher), 50 μg/mL streptomycin (Thermo Fisher), and 1 μg/mL amphotericin B (Thermo Fisher). Leaflets were transferred to a 250 units type II collagenase (Worthington) per mL EBSS solution and incubated at 37°C and 5% CO_2_ for 30 minutes under constant shaking. Leaflets were vortexed at maximum speed for 30 seconds, transferred to fresh collagenase solution, and incubated at 37°C and 5% CO_2_ for 60 minutes under constant shaking. Leaflets were vortexed again for 2 minutes at maximum speed and cells were passed through a 100 μm cell strainer using sterile transfer pipettes. Cells were centrifuged at 0.2*g* for 10 minutes, and pellets were resuspended in VIC expansion medium consisting of Media 199 (Life Technologies), 15% fetal bovine serum (FBS, Life Technologies), 50 U/mL penicillin, 50 μg/mL streptomycin, and 1 μg/mL amphotericin B. Cells were cultured at 37°C and 5% CO_2_ on tissue culture treated polystyrene (TCPS) for expansion before experiments. 70-80% confluent VIC cultures were harvested using trypsin (Life Technologies) and counted using an automated hemocytometer. VICs were seeded on PEG hydrogels at a density of 20,000 cells per cm^2^ growth area in Media 199 supplemented with 1% serum (FBS or human serum samples), 50 U/mL penicillin, 50 μg/mL streptomycin, and 1 μg/mL amphotericin B.

#### RT-PCR

RNA was extracted from male or female aortic valvular interstitial cells in culture using a RNeasy Micro Kit (Qiagen) according to the manufacturer’s protocol. RNA quality was assessed via spectrophotometry (ND-1000, NanoDrop), and cDNA was synthesized using an iScript Synthesis kit (Bio-Rad) according to the manufacturer’s protocol. Relative mRNA expression was determined using SYBR Green reagents on an iCycler (Bio-Rad). Normalizations were performed using the *RPL30* gene. Primer sequences are provided in **Table 1**.

**Table 1:**
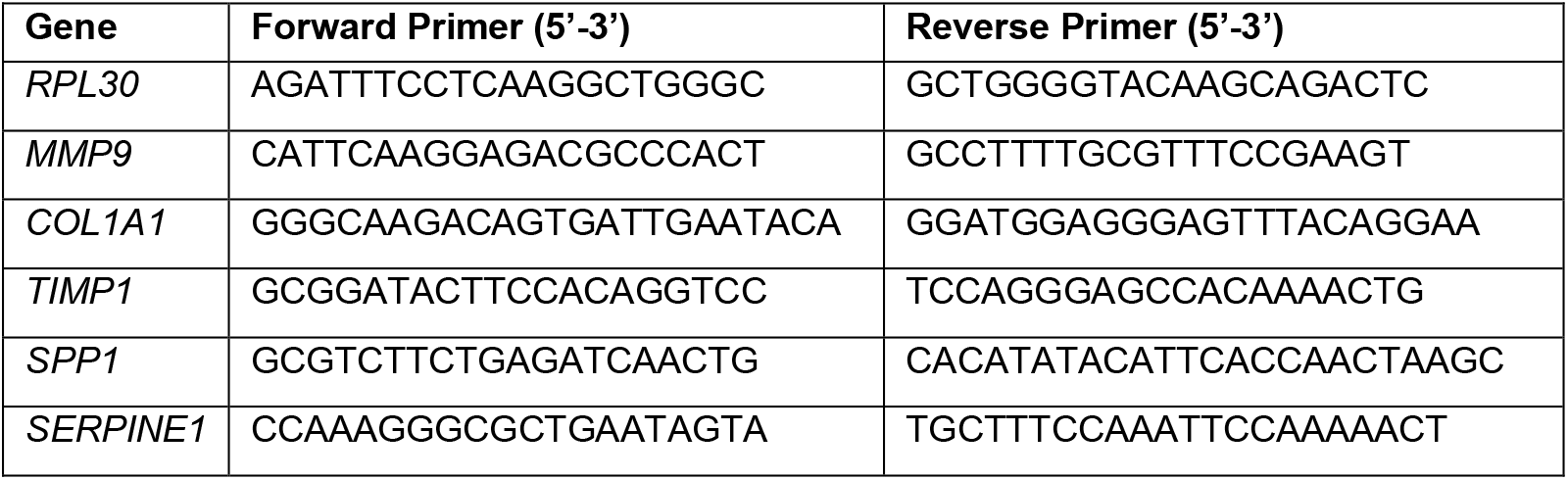
Primer sequences.

### Statistical analyses

Computational model data are shown as mean levels across the ensemble of model parameter sets. Significant differences between pre-TAVR and post-TAVR groups were determined using two-tailed Student’s t-tests in MATLAB. Correlations between activity data were determined using Pearson correlation in MATLAB, and Student’s t-tests with Benjamini-Hochberg correction for multiple comparisons were used to assess statistical significance. An aggregated matrix content score (MCS) was also derived for each patient from individual output activity levels using rank-normalized values. Model outputs were rank-normalized for each patient, and mean activity levels for all procollagens and matricellular proteins (Activity_Matrix_), mean activity levels for all matrix proteases (Activity_Protease_), and mean activity levels for all protease inhibitors were used for aggregation: *MCS* = *Activity_Matrix_* − *Activity*_*Protease*_ + *Activity*_*Inhib*_. Output nodes were categorized as follows: Activity_Matrix_: proCI, proCIII, fibronectin, periostin, osteopontin, LOXL1, P4H; Activity_Protease_: proMMPs 1, 2, 3, 8, 9, 12, 14, CTSC, CTSL; Activity_Inhib_: TIMP1, TIMP2, PAI1. Correlations involving MCS levels were determined using Spearman correlation in MATLAB and Student’s t-tests for statistical significance.

In vitro experimental mRNA measurements were analyzed using a one-way ANOVA with post-hoc Sidak’s post-hoc tests to directly compare bosentan treated vs. untreated groups in each patient-specific serum context. Six replicates were measured for each condition.

## References

1. Dweck, M. R., Boon, N. A. & Newby, D. E. Calcific aortic stenosis: A disease of the valve and the myocardium. J. Am. Coll. Cardiol. 60, 1854–1863 (2012).

2. Durko, A. P., Osnabrugge, R. L. & Kappetein, A. P. Long-term outlook for transcatheter aortic valve replacement. Trends Cardiovasc. Med. 28, 174–183 (2018).

3. Azevedo, C. F. et al. Prognostic Significance of Myocardial Fibrosis Quantification by Histopathology and Magnetic Resonance Imaging in Patients With Severe Aortic Valve Disease. J. Am. Coll. Cardiol. 56, 278–287 (2010).

4. Milano, A. D. et al. Prognostic value of myocardial fibrosis in patients with severe aortic valve stenosis. J. Thorac. Cardiovasc. Surg. 144, 830–837 (2012).

5. Yarbrough, W. M., Mukherjee, R., Ikonomidis, J. S., Zile, M. R. & Spinale, F. G. Myocardial remodeling with aortic stenosis and after aortic valve replacement: Mechanisms and future prognostic implications. J. Thorac. Cardiovasc. Surg. 143, 656–664 (2012).

6. Borer, J. S. & Sharma, A. Drug Therapy for Heart Valve Diseases. Circulation 132, 1038–1045 (2015).

7. Wang, H., Leinwand, L. A. & Anseth, K. S. Cardiac valve cells and their microenvironment-insights from in vitro studies. Nat. Rev. Cardiol. 11, 715–727 (2014).

8. Yip, C. Y. Y. & Simmons, C. A. The aortic valve microenvironment and its role in calcific aortic valve disease. Cardiovascular Pathology 20, 177–182 (2011).

9. Huang, H. Y. S., Liao, J. & Sacks, M. S. In-situ deformation of the aortic valve interstitial cell nucleus under diastolic loading. J. Biomech. Eng. 129, 880–889 (2007).

10. Rogers, J., Holmes, J., Saucerman, J. & Richardson, W. Mechano-Chemo Signaling Interactions Modulate Matrix Production by Cardiac Fibroblasts. bioRxiv 2020.05.06.077479 (2020). doi:10.1101/2020.05.06.077479

11. Davis, J. & Molkentin, J. D. Myofibroblasts: Trust your heart and let fate decide. J. Mol. Cell. Cardiol. 70, 9–18 (2014).

12. Aguado, B. A. et al. Transcatheter aortic valve replacements alter circulating serum factors to mediate myofibroblast deactivation. Sci. Transl. Med. 11, eaav3233 (2019).

13. Grim, J. C. et al. Secreted Factors From Proinflammatory Macrophages Promote an Osteoblast-Like Phenotype in Valvular Interstitial Cells. Arterioscler. Thromb. Vasc. Biol. 40, E296–E308 (2020).

14. Kural, M. H. & Billiar, K. L. Myofibroblast persistence with real-time changes in boundary stiffness. Acta Biomater. 32, 223–230 (2016).

15. Wang, H., Haeger, S. M., Kloxin, A. M., Leinwand, L. a & Anseth, K. S. Redirecting valvular myofibroblasts into dormant fibroblasts through light-mediated reduction in substrate modulus. PLoS One 7, e39969 (2012).

16. Kloxin, A. M., Benton, J. A. & Anseth, K. S. In situ elasticity modulation with dynamic substrates to direct cell phenotype. Biomaterials 31, 1–8 (2010).

17. Zeigler, A. C., Richardson, W. J., Holmes, J. W. & Saucerman, J. J. A computational model of cardiac fibroblast signaling predicts context-dependent drivers of myofibroblast differentiation. J. Mol. Cell. Cardiol. 94, 72–81 (2016).

18. Tan, P. M., Buchholz, K. S., Omens, J. H., McCulloch, A. D. & Saucerman, J. J. Predictive model identifies key network regulators of cardiomyocyte mechano-signaling. PLoS Comput. Biol. 13, 1–17 (2017).

19. Irons, L. & Humphrey, J. D. Cell signaling model for arterial mechanobiology. PLOS Comput. Biol. 16, e1008161 (2020).

20. Osmanbeyoglu, H. U., Pelossof, R., Bromberg, J. F. & Leslie, C. S. Linking signaling pathways to transcriptional programs in breast cancer. Genome Res. 24, 1869–1880 (2014).

21. Zeigler, A. C., Richardson, W. J., Holmes, J. W. & Saucerman, J. J. Computational modeling of cardiac fibroblasts and fibrosis. J. Mol. Cell. Cardiol. 93, 73–83 (2016).

22. Lacraz, G. P. A. et al. Tomo-Seq Identifies SOX9 as a Key Regulator of Cardiac Fibrosis during Ischemic Injury. Circulation 136, 1396–1409 (2017).

23. Scharf, G. M. et al. Inactivation of Sox9 in fibroblasts reduces cardiac fibrosis and inflammation. JCI Insight 4, (2019).

24. Jenke, A. et al. Transforming growth factor-β1 promotes fibrosis but attenuates calcification of valvular tissue applied as a three-dimensional calcific aortic valve disease model. Am. J. Physiol. Circ. Physiol. 319, H1123–H1141 (2020).

25. Sharma, V., Dogra, N., Saikia, U. N. & Khullar, M. Transcriptional regulation of endothelial-to-mesenchymal transition in cardiac fibrosis: Role of myocardin-related transcription factor A and activating transcription factor 3. Canadian Journal of Physiology and Pharmacology 95, 1263–1270 (2017).

26. Li, Y. et al. Cardiac Fibroblast–Specific Activating Transcription Factor 3 Protects Against Heart Failure by Suppressing MAP2K3-p38 Signaling. Circulation 135, 2041–2057 (2017).

27. Zhou, X. et al. Notch signaling inhibits cardiac fibroblast to myofibroblast transformation by antagonizing TGF-β1/Smad3 signaling. J. Cell. Physiol. 234, 8834–8845 (2019).

28. Kraeutler, M. J., Soltis, A. R. & Saucerman, J. J. Modeling cardiac β-adrenergic signaling with normalized-Hill differential equations: comparison with a biochemical model. BMC Syst. Biol. 4, 157 (2010).

29. Schroer, A. K., Ryzhova, L. M. & Merryman, W. D. Network Modeling Approach to Predict Myofibroblast Differentiation. Cell. Mol. Bioeng. 7, 446–459 (2014).

30. Morris, M. K., Saez-Rodriguez, J., Clarke, D. C., Sorger, P. K. & Lauffenburger, D. A. Training Signaling Pathway Maps to Biochemical Data with Constrained Fuzzy Logic: Quantitative Analysis of Liver Cell Responses to Inflammatory Stimuli. PLoS Comput. Biol. 7, e1001099 (2011).

31. Aguado, B., Grim, J., Rosales, A., Watson-Capps, J. & Anseth, K. Engineering precision biomaterials for personalized medicine. Sci. Transl. Med. 10, (2018).

32. Zeng, Q. et al. Notch1 Promotes the Pro-Osteogenic Response of Human Aortic Valve Interstitial Cells via Modulation of ERK1/2 and Nuclear Factor-κB Activation. Arterioscler. Thromb. Vasc. Biol. 33, 1580–1590 (2013).

33. Chen, J. et al. Notch1 Mutation Leads to Valvular Calcification Through Enhanced Myofibroblast Mechanotransduction. Arterioscler. Thromb. Vasc. Biol. 35, 1597–1605 (2015).

34. Menon, V. & Lincoln, J. The Genetic Regulation of Aortic Valve Development and Calcific Disease. Frontiers in Cardiovascular Medicine 5, 162 (2018).

35. Rutkovskiy, A. et al. Valve interstitial cells: The key to understanding the pathophysiology of heart valve calcification. J. Am. Heart Assoc. 6, 1–23 (2017).

36. Fischer, P. & Hilfiker-Kleiner, D. Survival pathways in hypertrophy and heart failure: The gp130-STAT3 axis. Basic Res. Cardiol. 102, 393–411 (2007).

37. Dziemidowicz, M. et al. The role of interleukin-6 in intracellular signal transduction after chronic β-adrenergic stimulation in mouse myocardium. Arch. Med. Sci. 15, 1565–1575 (2019).

38. Cucoranu, I. et al. NAD(P)H oxidase 4 mediates transforming growth factor-β1-induced differentiation of cardiac fibroblasts into myofibroblasts. Circ. Res. 97, 900–907 (2005).

39. Zhao, Q. D. et al. NADPH oxidase 4 induces cardiac fibrosis and hypertrophy through activating Akt/mTOR and NFκB signaling pathways. Circulation 131, 643–655 (2015).

40. Hagler, M. A. et al. TGF-β signalling and reactive oxygen species drive fibrosis and matrix remodelling in myxomatous mitral valves. Cardiovasc. Res. 99, 175–184 (2013).

41. Obana, M. et al. Therapeutic activation of signal transducer and activator of transcription 3 by interleukin-11 ameliorates cardiac fibrosis after myocardial infarction. Circulation 121, 684–691 (2010).

42. Su, S. A. et al. EphrinB2 regulates cardiac fibrosis through modulating the interaction of Stat3 and TGF-β/Smad3 signaling. Circ. Res. 121, 617–627 (2017).

43. Chakraborty, D. et al. Activation of STAT3 integrates common profibrotic pathways to promote fibroblast activation and tissue fibrosis. Nat. Commun. 8, (2017).

44. Deng, X. S., Meng, X., Song, R., Fullerton, D. & Jaggers, J. Rapamycin Decreases the Osteogenic Response in Aortic Valve Interstitial Cells Through the Stat3 Pathway. Ann. Thorac. Surg. 102, 1229–1238 (2016).

45. Seoudy, H. et al. Periprocedural Changes of NT-proBNP Are Associated With Survival After Transcatheter Aortic Valve Implantation. J. Am. Heart Assoc. 8, e010876 (2019).

46. Imai, K. et al. C-Reactive protein predicts severity, progression, and prognosis of asymptomatic aortic valve stenosis. Am. Heart J. 156, 713–718 (2008).

47. Huynh-Thu, V. A., Irrthum, A., Wehenkel, L. & Geurts, P. Inferring Regulatory Networks from Expression Data Using Tree-Based Methods. PLoS One 5, e12776 (2010).

48. Mercatelli, D., Scalambra, L., Triboli, L., Ray, F. & Giorgi, F. M. Gene regulatory network inference resources: A practical overview. Biochim. Biophys. Acta - Gene Regul. Mech. 1863, 194430 (2020).

49. Moerman, T. et al. GRNBoost2 and Arboreto: Efficient and scalable inference of gene regulatory networks. Bioinformatics 35, 2159–2161 (2019).

50. Rouillard, A. D. et al. The harmonizome: a collection of processed datasets gathered to serve and mine knowledge about genes and proteins. Database (Oxford). 2016, (2016).

51. Harris, C. R. et al. Array programming with NumPy. Nature 585, 357–362 (2020).

52. McKinney, W. Data Structures for Statistical Computing in Python. in Proceedings of the 9th Python in Science Conference (eds. van der Walt, S. J. & Millman, K. J.) 56–61 (2010). doi:10.25080/majora-92bf1922-00a

53. Shannon, P. et al. Cytoscape: A software Environment for integrated models of biomolecular interaction networks. Genome Res. 13, 2498–2504 (2003).

54. Zeigler, A. C. et al. Computational model predicts paracrine and intracellular drivers of fibroblast phenotype after myocardial infarction. Matrix Biol. 91–92, 136–151 (2020).

55. Mabry, K., Lawrence, R. & Anseth, K. Dynamic stiffening of poly(ethylene glycol)-based hydrogels to direct valvular interstitial cell phenotype in a three-dimensional environment. Biomaterials 49, 47–56 (2015).

