## Supplemental Figures for "Network Modeling Predicts Personalized Gene Expression and Drug Responses in Valve Myofibroblasts Cultured with Patient Sera"

### Supplementary Figures

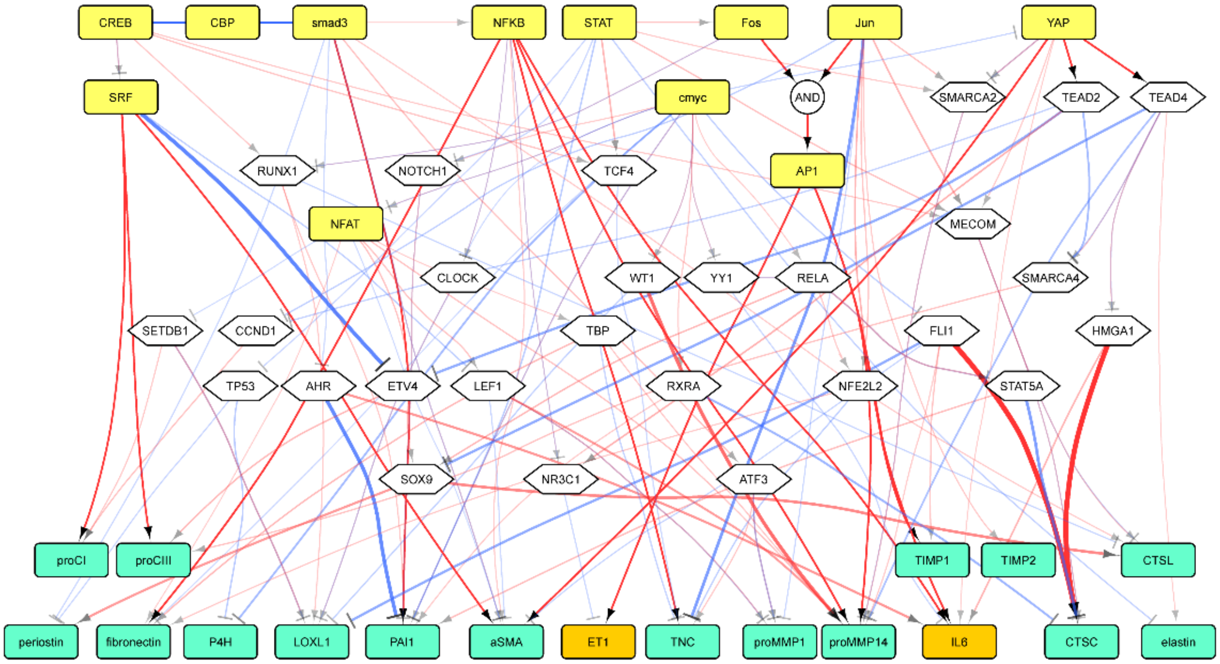


**Figure S1. Schematic of inferred transcriptional network.** Signaling-activated TFs (yellow boxes), secondary TFs (white hexagons), and model outputs associated with fibrosis or autocrine feedback (green/orange boxes) are connected by directed activation and inhibition reactions (red and blue arrows, respectively). Edge widths and transparency represent relative “importance” scores of each TF-target interaction as measured via the GRNBoost2 algorithm.

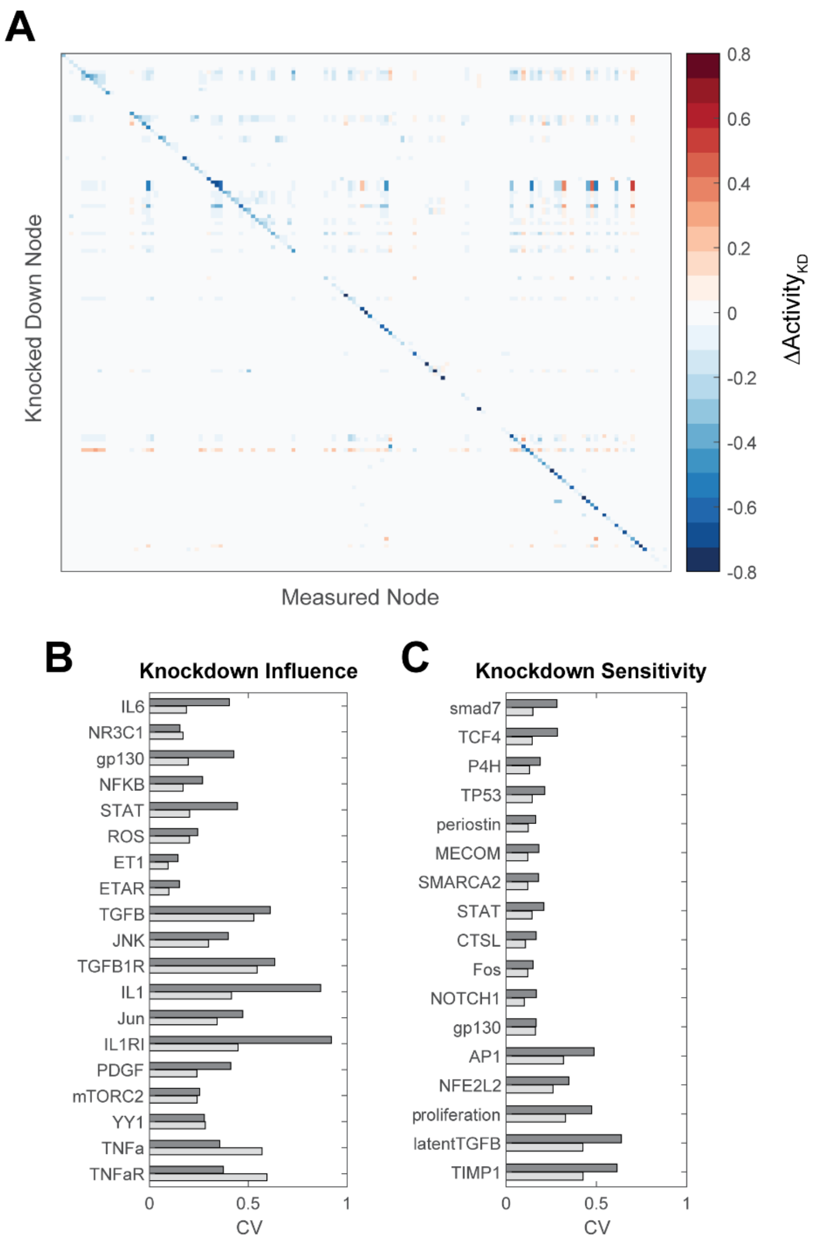


**Figure S2. Full network perturbation analysis results.** (A) Changes in activity for all nodes in the network were measured following comprehensive knockdown of individual nodes (Ymax = 0.1) under individual patient pre- and post-TAVR conditions. Values reflect average changes in node activity between perturbed and un-perturbed conditions with each patient condition. Refer to the model logic file available on GitHub for the order of nodes perturbed/measured. (B/C) Coefficients of variation (CV) across patient cohort for node influence and sensitivity values.

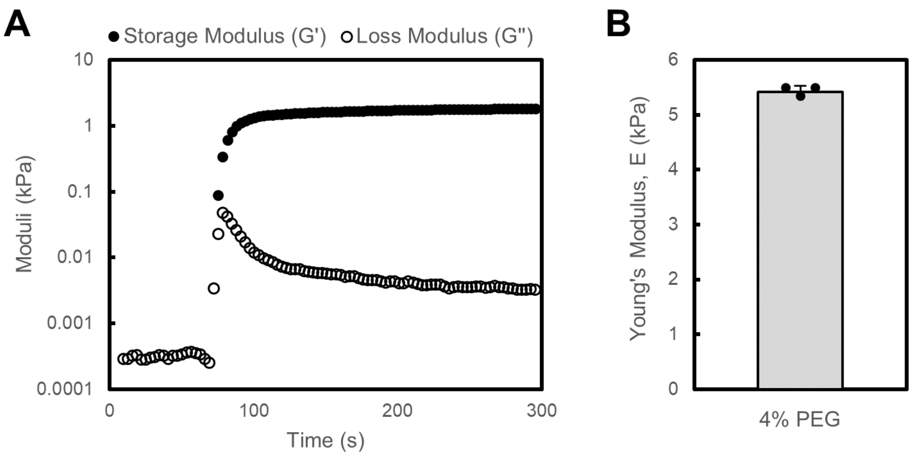


**Figure S3. Rheology analysis of PEG gel mechanics.** (A) Mechanical stiffness properties of 4% poly(ethylene glycol) (PEG) hydrogels were assessed during photopolymerization in situ within a parallel plate rheometer, which demonstrated typical increases in storage modulus and loss modulus during gelation. (B) Young's modulus calculated across 3 gels indicated highly consistent stiffness of the gels used for in vitro cell culture experiments.
